# Comparative Patterns of Variation on the X Chromosome and Autosomes: The Role of the Breeding Sex Ratio

**DOI:** 10.1101/2025.01.17.633266

**Authors:** William J. Spurley, Bret A. Payseur

## Abstract

In many populations, unequal numbers of females and males reproduce each generation. This imbalance in the breeding sex ratio (BSR) shapes patterns of genetic variation on the sex chromosomes and the autosomes in distinct ways. Despite recognition of this phenomenon, effects of the BSR on some aspects of variation remain unclear, especially for populations with non-equilibrium demographic histories. To address this gap in the field, we used coalescent simulations to examine relative patterns of variation at X-linked loci and autosomal loci in populations spanning the range of BSR with historical changes in population size. Shifts in BSR away from 1:1 reduce nucleotide diversity and the number of unique haplotypes and increase linkage disequilibrium and the frequency of the most common haplotype, with contrasting effects on X-linked loci and autosomal loci. Strong population bottlenecks transform relationships between the BSR, the site frequency spectrum, and linkage disequilibrium while relationships between the BSR, nucleotide diversity, and haplotype characteristics are broadly conserved. Our findings indicate that evolutionary interpretations of variation on the X chromosome should consider the combined effects of the BSR and demographic history. The genomic signatures we report could be used to reconstruct these fundamental population parameters from genomic data in natural populations.

**Significance Statement:** The breeding sex ratio is a fundamental evolutionary parameter, but genomic analyses routinely assume it is 1:1. Our research characterizes the relationships between the breeding sex ratio and multiple facets of genomic variation and shows how these relationships change in the context of dynamic demographic histories. In doing so, we provide increasingly realistic expectations for patterns of X-linked and autosomal variation in population genomic datasets collected from natural populations.

## Introduction

Dioecious organisms have two biological parents. Despite this fact, females and males can contribute unequally to the next generation from an evolutionary perspective. Polygyny (the mating of males with multiple females) biases the breeding sex ratio (BSR) away from 1:1 toward females, whereas polyandry (the mating of females with multiple males) pushes the BSR toward males. Even in populations in which most members mate monogamously, extra-pair mating can distort the BSR away from 1:1 (Fernandez-Duque and Huck 2013; Huck et al. 2020). Sexual dimorphism in life-history traits, including generation time and lifespan, can also lead to unequal reproductive contributions by females and males (Lawler 2009; Amster and Sella 2020). Consequently, populations with BSRs that depart from 1:1 are expected to be common in nature.

Regardless of the cause of shifts in the BSR, this fundamental demographic parameter is predicted to leave signatures in genomic patterns of variation. The BSR shapes the effective population size (N_e_), which captures the effects of genetic drift and is a key determinant of genetic variation (Wright 1940; Kimura and Crow 1963; Waples 2022). At autosomal loci, N_e_ is reduced as the BSR moves away from 1:1 (Hill 1972; Crow and Denniston 1988). The decline in N_e_ is symmetrical because a male bias (a higher proportion of breeding males in the population) and a female bias (a higher proportion of breeding females in the population) both decrease the number of unique autosomes contributing to the next generation (Caballero 1995).

The effects of the BSR on sex-linked loci are distinct from those on autosomal loci because the sex chromosomes are inherited differently. The N_e_ of non-recombining Y-linked loci reflects the number of unique breeding males in the population. When the BSR is 1:1 in a population of constant size, the N_e_ of Y-linked loci is expected to be ¼ the N_e_ of autosomal loci (Caballero 1995). A female bias in the BSR shifts the ratio of N_e_ for Y-linked loci and autosomal loci below ¼ while a male bias moves the ratio above ¼ (Wright 1933). The relationship between X-linked loci and the BSR is more complex. Due to hemizygosity of the X chromosome in males, the N_e_ of X-linked loci is expected to be ¾ the N_e_ of autosomal loci in a population of constant size (Wright 1933; Hill 1972). When the BSR is biased toward males, there are fewer total copies of the X chromosome in the breeding population, and the N_e_ of X-linked loci is lower than ¾ the N_e_ of autosomal loci (Caballero 1995). As the BSR shifts away from 1:1 in favor of females, the N_e_ for X-linked loci becomes more similar to the N_e_ of autosomal loci, and even exceeds it at extreme female biases (Caballero 1995; Musharoff 2019). Expectations are reversed for species with Z/W sex chromosomes, where females are the heterogametic sex.

The effects of the BSR on N_e_ provide a straightforward path to understanding the connection between the BSR and patterns of genetic variation. The population mutation rate (θ), a primary determinant of levels of genetic variation under neutrality, is a product of N_e_ and the per-site, per-generation mutation rate (Watterson 1975). The population recombination rate (ρ), a primary determinant of linkage disequilibrium (LD) and haplotype structure, is a product of N_e_ and the per-site, per-generation recombination rate (Hill 1981; Hudson 1987). Changes in the BSR alter these parameters in different ways for X-linked loci and autosomal loci. For example, a male bias disproportionately reduces θ at X-linked loci relative to autosomal loci, with concomitant effects on nucleotide diversity (Charlesworth 2001). A male bias differentially inflates LD at X-linked loci compared to autosomal loci because two factors decrease ρ: reduced N_e_ and a lack of recombination in males (Labuda et al. 2010; Lohmueller et al. 2010). The disparate effects of the BSR on patterns of genetic variation at X-linked loci and autosomal loci enable departures from a 1:1 BSR to be detected from population genomic data. Nucleotide diversity, the site frequency spectrum (SFS), and LD point to female biases in human populations (Hammer et al. 2008; Emery et al. 2010; Musharoff et. 2019; Labuda et al. 2010).

Patterns of variation at X-linked loci and autosomal loci can also deviate from neutral expectations due to demographic history. A reduction in population size initially decreases X-linked diversity more substantially than autosomal diversity (Pool and Nielsen, 2007). However, when the population size increases again, as in the case of a population bottleneck, X-linked diversity recovers to its original level faster than autosomal diversity (Pool and Nielsen 2007). Established effects of demographic history on the SFS (Tajima 1989; Griffiths and Tavaré 1998; Gutenkunst et al. 2009), LD (Waples 2006; Santiago et al. 2020), and haplotype structure (Lohmueller et al. 2009; Palamara et al. 2012) raise the prospect that these metrics could also differ between X-linked loci and autosomal loci when population size fluctuates (Wall et al. 2002).

Although departures from a 1:1 BSR are expected to be common in natural populations, gaps persist in our understanding of how this fundamental reproductive parameter shapes patterns of genetic variation. Importantly, little is known about how the BSR and demographic history *jointly* affect X-linked diversity and autosomal diversity (Webster and Wilson Sayres 2016; Musharoff et al. 2019). Moreover, theoretical and empirical studies connecting the BSR to genetic variation have focused on levels of diversity (Hammer et al. 2010; Emery et al. 2010; Clemente et al. 2018; Musharoff et al. 2019); expectations for measures of LD and haplotype structure remain unclear. To address these challenges, we examine the role of BSR in shaping variation at X-linked loci and autosomal loci in the context of realistic features of demographic history.

## Results

To understand the joint effects of the BSR and demographic history on genomic patterns of sequence variation, we performed independent coalescent simulations of 1,000 unlinked loci on the X chromosome and 1,000 unlinked loci on the autosomes. We measured patterns of variation using a suite of common summary statistics that collectively capture levels of diversity, the site frequency spectrum (SFS) of polymorphisms, haplotype structure, and linkage disequilibrium (LD). To illustrate how the BSR interacts with characteristics of demographic history to distort patterns of variation observed at mutation-recombination-drift equilibrium, we examined population bottlenecks with three different strengths, durations, and timings across the spectrum of BSR. Results for a wider range of scenarios are presented in the Supplementary Material.

### Effects of the BSR on Patterns of Variation at Mutation-Recombination-Drift Equilibrium

To first establish how the BSR alone affects variation on X-linked loci and autosomal loci, we focused on sex biases in the absence of demographic change. Some of the patterns we describe in this section have been noted previously; we reintroduce them here to provide immediate comparison to scenarios in which both sex biases and demographic change are present (and to validate our simulation framework).

Average nucleotide diversity (π, Figure 1A) across autosomal loci and across X-linked loci recaptures established trends (Charlesworth 2001; Musharoff et al. 2019). For autosomal loci, π declines symmetrically with distance from a maximum at a proportion of females in the breeding population (pf) = 0.5 (a 1:1 BSR). Alternatively, for X-linked loci, π is maximized with a slight female bias and declines similarly to the autosomal scenario as the BSR shifts away from this point. X-linked and autosomal diversity are similar at more extreme sex biases, particularly with a female bias. These patterns apply equally to Watterson’s θ (Figure 1B), resulting in no change in Tajima’s D (which is computed as the normalized difference between π and Watterson’s θ) across all BSRs (Figure 1C).

**Figure 1.**
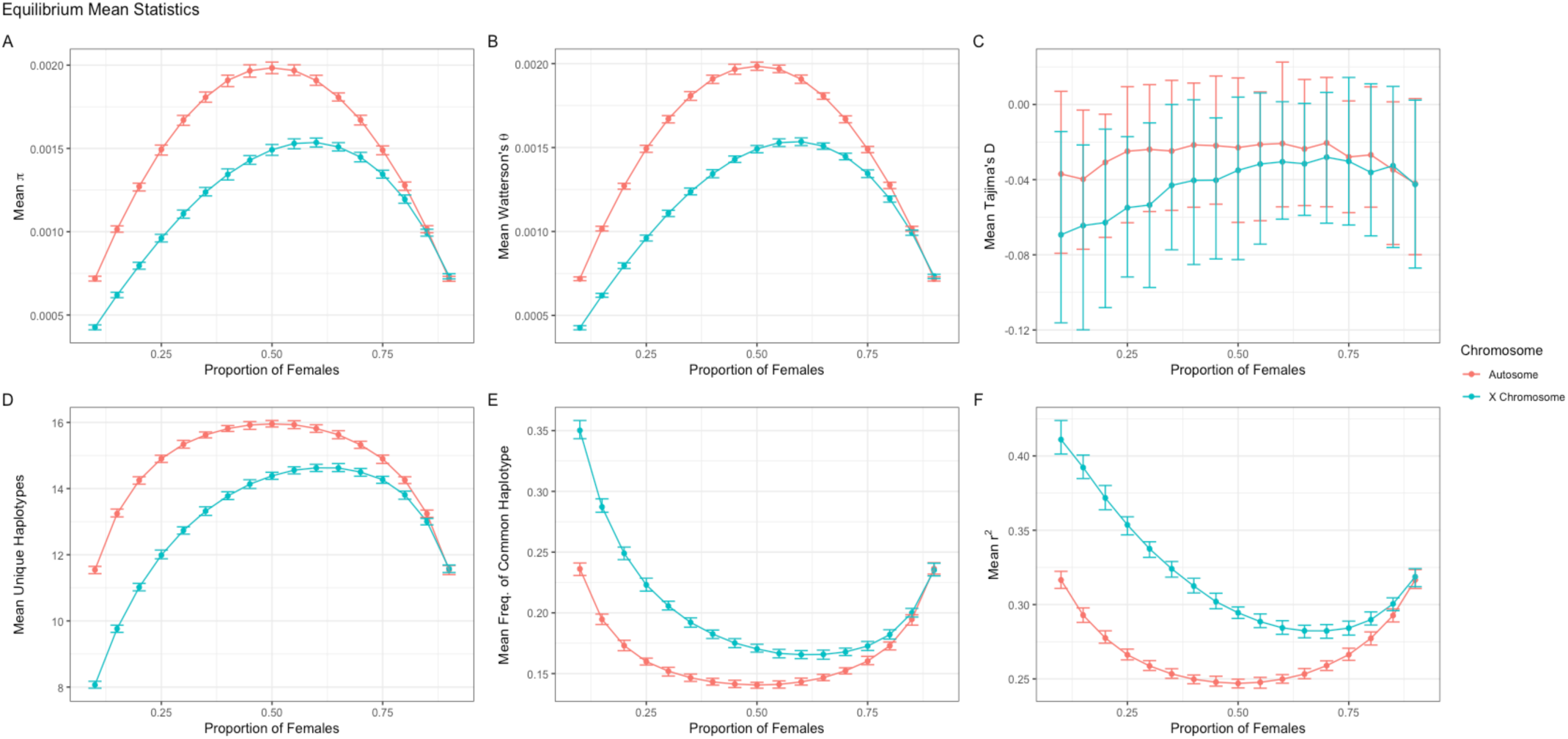
Patterns of variation are sensitive to the BSR. Mean π (A), Watterson’s θ (B), Tajima’s D (C), number of haplotypes (D), frequency of the common haplotype (E), and r^2^ (F) under a constant population size at varying breeding sex ratios. Points represent mean values across 100 evolutionary replicates of 1000 independent loci for X-linked loci (blue) and autosomal loci (red). Error bars summarize the 95% confidence interval by taking the 97.5^th^ and 2.5^th^ quantile of each distribution of replicates.

Effects of the BSR extend to summary statistics that capture haplotypes and LD. The average number of unique haplotypes across autosomal loci declines with stronger sex biases in each direction (Figure 1D). For X-linked loci, the average number of unique haplotypes is maximized at a slight female bias. Although these trends resemble patterns for π and Watterson’s θ, the number of haplotypes decays more slowly than π as the BSR moves away from 0.5. For X-linked loci, the frequency of the most common haplotype is maximized at extreme male bias, when the number of haplotypes is minimized (Figure 1E). Autosomal loci show higher r^2^ at extreme BSRs (Figure 1F). For X-linked loci, r^2^ is maximized at an extreme male bias and minimized at a slight female bias.

The inter-locus variance of each summary statistic is also responsive to changes in the BSR (Supplementary Figure 1). The variance of π and the variance of Watterson’s θ follow patterns for the averages of these statistics described above. In contrast to average Tajima’s D, the variance of Tajima’s D is affected by changes in the BSR, with elevated variance at stronger sex biases. The frequency of the most common haplotype and r^2^ also follow this pattern, but with less change across moderate BSR values. The variance in the number of haplotypes generally increases with the degree of sex bias, except for X-linked loci at the strongest male bias (pf = 0.1).

### Effects of the BSR and Bottleneck Strength on Patterns of Variation

To determine how the BSR interacts with demographic history to shape patterns of variation, we examined three different bottleneck strengths: a 50% reduction, an 80% reduction, and a 95% reduction from the ancestral size. For results reported in this section, we held duration and timing constant: bottlenecks lasted 500 generations and ended 500 generations ago.

Under a bottleneck, the relationships between the BSR and average levels of variation (π and Watterson’s θ) for X-linked loci and autosomal loci resemble those in constant-size populations (Figure 2A and 2B). For stronger bottlenecks, Watterson’s θ is reduced more than π, particularly at moderate sex biases, which induces a dependence of Tajima’s D on the BSR (Figure 2C). At an 80% reduction, average Tajima’s D begins to differentiate at strong male biases, but not at any other BSR. At a 95% reduction, Tajima’s D is fully sensitive to the BSR: extreme sex biases confer negative Tajima’s D values, whereas moderate sex biases yield positive values.

**Figure 2.**
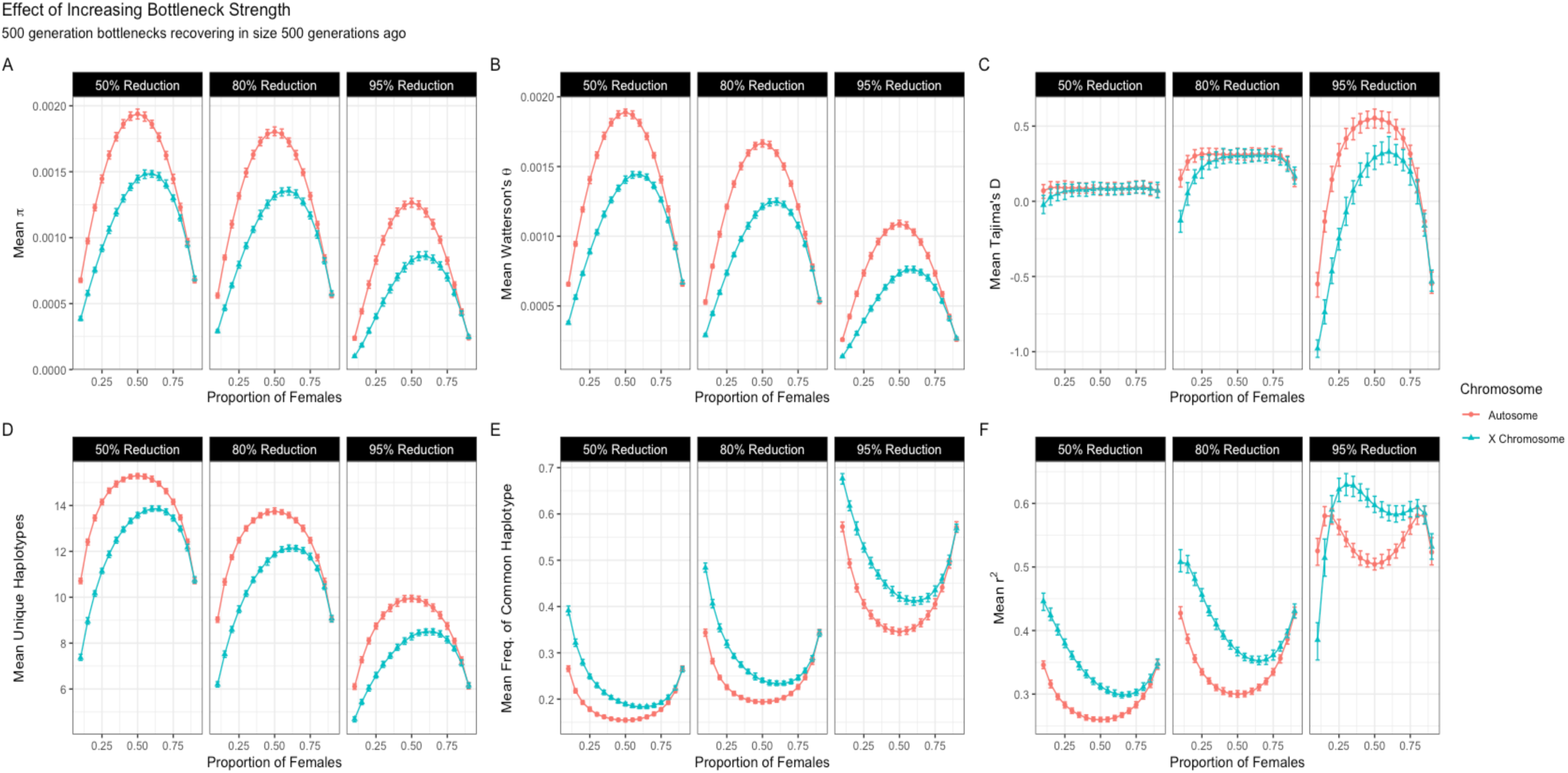
Increasing population bottleneck strength changes the relationship between the BSR and patterns of variation. Mean π (A), Watterson’s θ (B), Tajima’s D (C), number of haplotypes (D), frequency of the common haplotype (E), and r^2^ (F) under bottlenecks with variable strength, lasting 500 generations, and recovering in size 500 generations ago. Points represent mean values across 100 evolutionary replicates of 1000 independent loci for X-linked loci (blue) and autosomal loci (red). Error bars summarize the 95% confidence interval by taking the 97.5^th^ and 2.5^th^ quantile of each distribution of replicates. Each strength represents 50%, 80%, or 95% reduction from the ancestral population size (left, center, and right panels, respectively).

As bottleneck strength increases, the number of unique haplotypes declines and the frequency of the most common haplotype increases (Figure 2D and 2E). Still, relationships between the BSR and haplotype patterns with a bottleneck resemble those in populations of constant size.

Linkage disequilibrium shows a distinct pattern. As bottlenecks increase in strength, average r^2^ changes with the BSR in ways that depart from equilibrium patterns (Figure 2F). With 50% and 80% reductions in population size, r^2^ is elevated across most of the range of BSR, compared to equilibrium values. In contrast, a 95% reduction in population size transforms the relationship of the BSR and LD. In a surprising reversal of equilibrium patterns, r^2^ is higher at autosomal loci than at X-linked loci when male bias is extreme. LD is also maximized for both X-linked loci and autosomal loci at moderate sex biases.

We considered the effects of bottleneck strength across additional scenarios of bottleneck timing and duration. Our results consistently indicate that changing bottleneck strength has no strong effect on relative patterns of X-linked loci and autosomal loci for π, Watterson’s θ, and the haplotype summaries, regardless of bottleneck timing and duration. In contrast, bottlenecks shape the relationships between the BSR, Tajima’s D, and r^2^ in a manner that depends on each bottleneck parameter (Supplementary Figure 2 and 3). These findings demonstrate strong interactions between BSR and demographic history in determining the SFS and LD.

Increasing the strength of a bottleneck does not affect the relationship of the BSR with the inter-locus variances of π or Watterson’s θ and has little effect on the variances of r2 and the frequency of the common haplotype. (Supplementary Figure 4). In contrast, a 95% reduction in population size transforms the relationship between the variance of Tajima’s D and the BSR, leaving a signature resembling mean r^2^ at an 80% reduction (described above). The variance of the number of haplotypes is also sensitive to changes in bottleneck strength. With a weak bottleneck, the variance is greater for X-linked loci than autosomal loci (except when the male bias is extreme). As the strength of the bottleneck increases, the patterns of variance change among the moderate BSRs, leading to a higher variance in the haplotype count for autosomal loci than X-linked loci at a 95% reduction.

### Effects of the BSR and Bottleneck Duration on Patterns of Variation

To explore how bottleneck duration affects relationships between BSR and patterns of variation, we simulated 100-generation, 500-generation, and 1000-generation bottlenecks with a 95% reduction in size and a recovery 500 generations ago.

As bottleneck duration increases, average diversity is reduced, with relationships between the BSR and levels of variation mostly being maintained (Figure 3A and 3B). These effects are only observed with strong bottlenecks; weaker bottlenecks show little change in variation as duration increases (Supplementary Figure 5 and 6). Increasing bottleneck duration induces a strong dependence of BSR on Tajima’s D with bottlenecks lasting 500 or 1000 generations (Figure 3C). With a 100-generation bottleneck, there is little differentiation of average Tajima’s D at X-linked and autosomal loci except at an extreme male bias, where X-linked loci have a lower average value. For a 500-generation bottleneck, parabolic relationships emerge between the BSR and Tajima’s D. For autosomal loci, Tajima’s D is positive for all but the most extreme BSR values, whereas for X-linked loci, Tajima’s D is negative at a wider range of BSRs. When the bottleneck lasts for 1000 generations, Tajima’s D for both X-linked and autosomal loci is shifted to be more negative.

**Figure 3.**
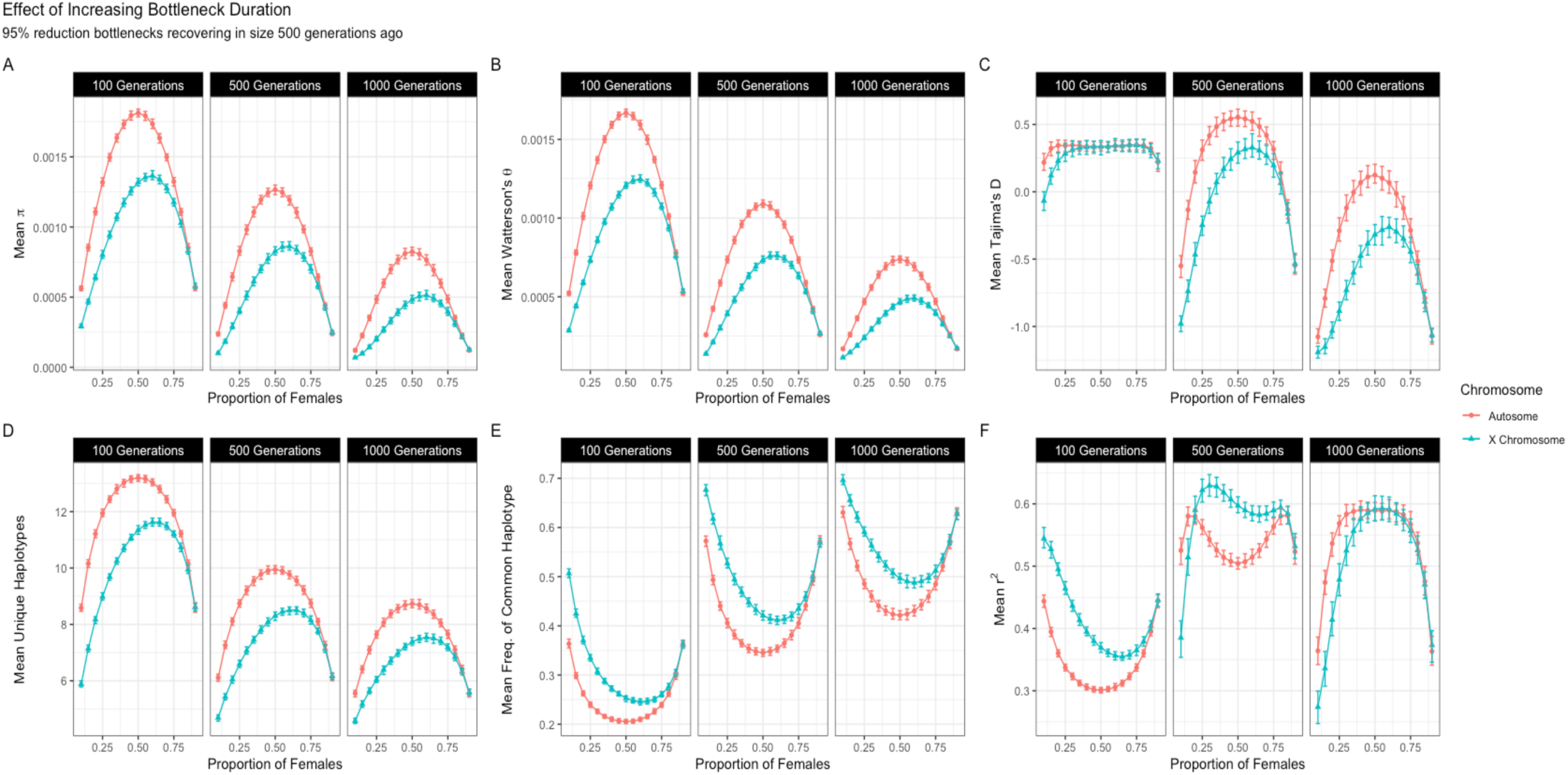
Increasing bottleneck duration changes the relationship between the BSR and patterns of variation. Mean π (A), Watterson’s θ (B), Tajima’s D (C), number of haplotypes (D), frequency of the common haplotype (E), and r^2^ (F) under bottlenecks with variable duration, a 95% reduction, and recovering in size 500 generations ago. Points represent mean values across 100 evolutionary replicates of 1000 independent loci for X-linked loci (blue) and autosomal loci (red). Error bars summarize the 95% confidence interval by taking the 97.5^th^ and 2.5^th^ quantile of each distribution of replicates. Each duration represents 100, 500, and 1000 generation long bottlenecks (left, center, and right panels, respectively).

Effects of the BSR on average haplotype number and average frequency of the most common haplotype do not change with bottleneck duration (Figure 3D and 3E). In contrast, LD shows a different relationship to the BSR at each bottleneck duration (Figure 3F). A 100-generation bottleneck leads to trends like those in the absence of a bottleneck, with average r^2^ elevated throughout. A 500-generation bottleneck induces complex relationships between the BSR and r^2^ that differ for strong and moderate sex biases (resembling the effects of a strong bottleneck, described above). At a 1000-generation bottleneck, there is little differentiation between X-linked loci and autosomal loci at a female bias. For both classes of loci, a stronger sex bias confers reduced r^2^ values, a stark contrast to the equilibrium scenario.

Relationships between the BSR and the inter-locus variances of π and Watterson’s θ are maintained as the duration of bottlenecks increases (Supplementary Figure 7). Nevertheless, the disparity between X-linked loci and autosomal loci is larger for longer bottlenecks. The variance of Tajima’s D again takes on patterns resembling those for mean r^2^ (see Figure 3F). For both r^2^ and the frequency of the most common haplotype, longer bottlenecks yield higher variances across the range of BSR, but there is a slight decline at the most extreme male bias. Variances in haplotype number are lower with extreme sex biases than at a balanced BSR.

### Effects of the BSR and Bottleneck Timing on Patterns of Variation

To examine how changes in bottleneck timing affect the relationships between the BSR and patterns of variation, we simulated bottlenecks that recover to the original population size 500, 1000, or 5000 generations ago while holding constant the strength (95% reduction) and duration (500 generations).

More recent bottlenecks result in lower average diversity across the range of BSRs (Figure 4A and 4B). With a bottleneck recovery 5000 generations ago, there is little change in average Tajima’s D with the BSR. In contrast, Tajima’s D is dependent on the BSR when bottlenecks end 1000 or 500 generations ago. A bottleneck recovering 500 generations ago yields more positive Tajima’s D than a distant bottleneck at a moderate sex bias (Figure 4C). Haplotype summaries maintain their relationships with the BSR regardless of the timing of bottlenecks (Figure 4D and 4E).

**Figure 4.**
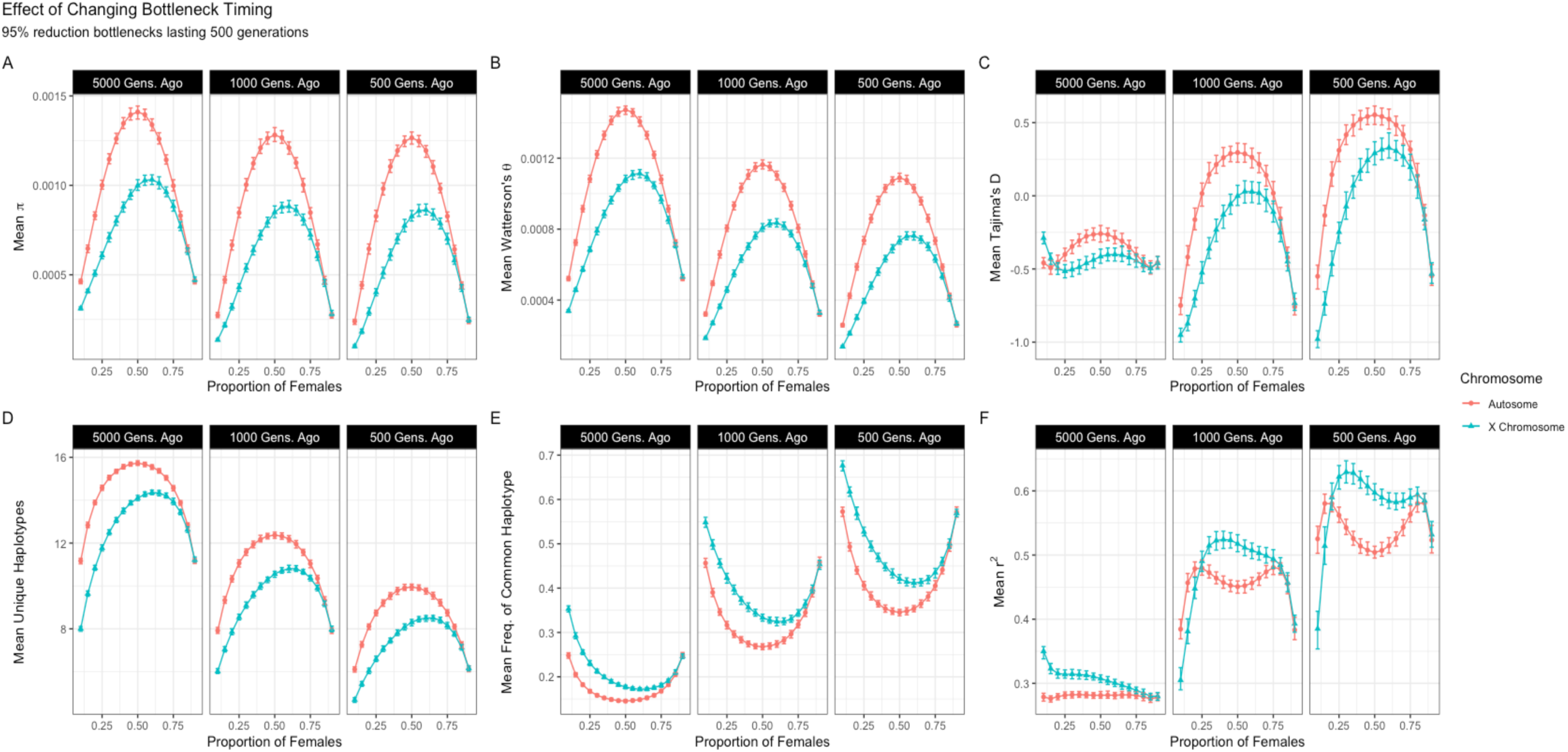
The timing of a bottleneck influences the relationship between the BSR and patterns of variation. Mean π (A), Watterson’s θ (B), Tajima’s D (C), number of haplotypes (D), frequency of the common haplotype (E), and r^2^ (F) under bottlenecks with variable timing, a 95% reduction, and lasting 500 generations. Points represent mean values across 100 evolutionary replicates of 1000 independent loci for X-linked loci (blue) and autosomal loci (red). Error bars summarize the 95% confidence interval by taking the 97.5^th^ and 2.5^th^ quantile of each distribution of replicates. Each timing represents bottlenecks recovering to the ancestral population size 5000, 1000, and 500 generations ago (left, center, and right panels, respectively).

Varying bottleneck timing alters the relationship between the BSR and LD (Figure 4F). When a bottleneck ends 5000 generations ago, average r^2^ at autosomal loci is largely unchanged across the range of BSRs. For X-linked loci, average r^2^ is maximized at an extreme male bias and minimized at an extreme female bias. When a bottleneck ends 1000 generations ago, autosomal r^2^ is maximized at two points: a strong male bias (pf = 0.2) and a strong female bias (pf = 0.8). For X-linked loci, average r^2^ increases from an extreme male bias to a moderate one (pf = 0.1 – 0.3), then among the moderate sex biases (pf = 0.3 – 0.8) it decreases gradually, before rapidly decreasing at a strong female bias (pf = 0.8 – 0.9). Similar patterns emerge for a bottleneck ending 500 generations ago, but with a greater difference between X-linked loci and autosomal loci at moderate sex biases. The ability to distinguish the effect of changes in bottleneck timing on patterns of variation is reduced for weaker bottlenecks (Supplementary Figure 8).

Although the timing of bottlenecks has little effect on the inter-locus variance of π and the variance of Watterson’s θ, the variance of Tajima’s D responds to changes in timing in patterns that resemble the average r^2^ (Supplementary Figure 9). The variance of the number of haplotypes is more sensitive to changes in bottleneck timing at moderate sex biases compared to extreme sex biases. For the frequency of the most common haplotype and r^2^, more recent bottlenecks lead to elevated variances, but less so for an extreme male bias.

### Investigating Patterns Sensitive to the Interaction between BSR and Demographic History

To further investigate interactions between the BSR and demographic history, we examined the SFS and LD in greater detail. First, we compared the full SFS (of which Tajima’s D is a summary) for X-linked loci and autosomal loci in a population with a severe bottleneck (95% strength, lasting 500 generations, and recovering 500 generations ago) and in a population of constant size, each across a range of BSR values.

Without changes in population size, the average SFS is the same in populations with different BSR values, as the proportion of SNPs in each frequency bin is independent of N_e_ (Fu, 1995; Supplementary Figure 10). With a bottleneck but no sex bias (pf = 0.5), the proportion of alleles with intermediate frequencies increases, but more noticeably for the autosomes (Supplementary Figure 11). This pattern aligns with the positive values for Tajima’s D observed for this demographic history and BSR (see Figure 2C, middle panel). With a bottleneck, the proportion of singletons increases with either male bias or female bias. This increase, which causes Tajima’s D to be more negative, marks a shift in abundance toward more recent mutations. With the loss of diversity caused by reductions in N_e_ from both a strong sex bias and a bottleneck a larger share of variants emerge after the recovery to the ancestral population size. As a result, the shape of the SFS and negative values of Tajima’s D reflect a population expansion.

To understand how LD at individual loci shape the average patterns of r^2^ described above, we explored the distribution of r^2^ across 1000 loci in a population with an extreme bottleneck (95% reduction in size, lasting 500 generations, and recovering 500 generations ago). Under this scenario, we observe an unexpected decline in average r^2^ at a strong male bias (Figure 4C, middle panel). Examining the distribution of r^2^ for X-linked loci at this strong male bias (pf = 0.1), we observe two distinct peaks around 0.1 and 0.9 (Supplementary Figure 12, top right panel). This pattern suggests that the reduced r^2^ at a strong male bias is driven by an increased share of loci with very little LD. One key difference between loci with r^2^ = 0.1 and loci with r^2^ = 0.9 is the time to the most recent common ancestor (TMRCA). Inspection of simulated genealogies reveals that loci with r^2^ < 0.1 have an average TMRCA of 906 generations while loci with r^2^ > 0.9 have an average TMRCA of 5090 generations (Supplementary Figure 12, bottom right panels). Whereas the average TMRCA for loci with an r^2^ < 0.1 falls within the epoch of reduced population size in this strong bottleneck, the average TMRCA for loci with an r^2^ > 0.9 is much older than the size change. Therefore, with a strong male bias and a strong bottleneck, average LD is reduced by an increase in the share of loci that experience faster coalescence and an altered demographic history.

### Effects of the BSR and Sex Differences in Mutation Rate on Patterns of Variation

To explore the capacity for sex-biased mutation to jointly shape patterns of variation with BSR and demographic history, we simulated the same scenarios described above, but with a male mutation rate three times the female mutation rate (Supplementary Figure 13). As expected, patterns of variation at autosomal loci are unaffected by this change. Additionally, r^2^ is unaltered for both X-linked loci and autosomal loci. At X-linked loci, π, Watterson’s θ, and the number of unique haplotypes are reduced by male-biased mutation, and the frequency of the most common haplotype increases. Effects are less severe for the two haplotype summaries than for π and Watterson’s θ. Similar patterns are observed with population bottlenecks, suggesting little interaction between sex differences in mutation rate and demographic history (Supplementary Figure 14).

## Discussion

From a genetic perspective, the X chromosome is special. Unequal transmission between the sexes and alternating ploidy between haploid males and diploid females makes it difficult to interpret patterns of variation on the X chromosome compared to autosomes (Hudson and Turelli 2003; Kirkpatrick et al. 2010). Partly because of these complications, the X chromosome is often ignored in population genomic studies. However, the biological factors that make the X chromosome difficult to compare with the autosomes also predict unique evolutionary dynamics. For example, the X chromosome offers the capacity to reveal differences between the sexes in the operation of evolutionary processes, including natural selection (Corl and Ellegren 2012) and migration (Goldberg and Rosenberg 2015).

The BSR must be considered when drawing evolutionary inferences from relative patterns of variation on the X chromosome and the autosomes. Our results confirm and extend this notion in two principal ways, illustrating the genomic consequences of the BSR for empirical datasets. First, we characterize the effects of the BSR across multiple axes of variation, providing a fuller picture of the genomic signatures of this fundamental reproductive parameter. Second, we distinguish facets of variation shaped by interactions between the BSR and demographic history (LD and SFS) from those for which relationships with the BSR are more stable (nucleotide diversity and haplotype characteristics).

Our conclusions are tempered by limitations of our study design. Our coalescent approach modeled the BSR and demographic parameters through their effects on N_e_ rather than modeling a specific mating structure. While this strategy enabled rapid and replicable simulations across the BSR and multiple demographic histories, additional insights could be gained through forward simulations that directly consider differences in inheritance between the X chromosome and the autosomes. We also assumed that all loci evolve neutrally. We sought to mimic a population genomic study that prioritizes intergenic loci far from genes for demographic inference, but isolating loci that evolve neutrally can be a challenge. Future efforts to understand effects of the BSR could consider linked selection (Maynard Smith and Haigh 1974; Charlesworth et al. 1993). Finally, we stipulated constancy of the BSR throughout population history, even as we might expect this parameter to shift over time due to changes in mating behavior and life-history traits.

Despite these caveats, our findings hold important implications for demographic inferences drawn from genomic patterns of variation. The SFS is commonly used to reconstruct historical population sizes (e.g. Gutenkunst et al. 2009), but estimates may be biased by unmodeled departures from a 1:1 BSR. As an instructive example, consider the SFS in a population with a 95% size reduction lasting 1000 generations that recovers in size 500 generations ago (Figure 4C, right panel). If the BSR is sufficiently female biased (pf > 0.6), Tajima’s D for autosomal loci is more negative than what is expected at a balanced BSR. The higher proportion of singletons driving this pattern could be misconstrued as solely due to demographic history if the BSR is not considered. Our results suggest that the mischaracterization of demographic history is more likely when populations experience strong bottlenecks. To minimize these complications, we encourage the continued development of methods with the capacity for joint estimation of the BSR and demographic history (Clemente et al. 2018; Musharoff et al. 2019).

Our results are also relevant to efforts to detect selection targeting the X chromosome (Begun and Whitley 2000; Kauer et al. 2002; Payseur and Nachman 2002; Gottipati et al. 2011; Nam et al. 2015; Harris and Garud 2023; Harris et al. 2024). Beneficial mutations that arise on the X chromosome enjoy shorter sojourn times, regardless of dominance (Avery 1984; Aquadro et al. 1994). Recessive, X-linked beneficial mutations also fix faster than their autosomal counterparts (Charlesworth et al. 1987), and the dominance conditions for faster X evolution are less restrictive when the BSR is female-biased (Laporte and Charlesworth 2002; Vicoso and Charlesworth 2009). Together, these factors predict reduced X-linked diversity compared to autosomal diversity (Begun and Whitley 2000; Betancourt et al. 2004). In contrast, background selection (Charlesworth et al. 1993) predicts higher than expected diversity on the X chromosome because hemizygosity exposes recessive, deleterious variants to selection at lower allele frequencies (Aquadro et al. 1994). Regardless of the selection being characterized, our results underscore the importance of incorporating the BSR alongside demographic history into null models used to characterize selection on the X chromosome from levels of diversity, the site frequency spectrum, linkage disequilibrium, and haplotype patterns (Veeramah et al. 2014; Harris et al. 2024).

Knowledge of the BSR can improve inferences about demographic history and selection, and comparisons between the X chromosome and autosomes make it possible to reconstruct this parameter from population genomic data. Building on earlier rejections of a balanced BSR in humans based on individual summary statistics (Hammer et al. 2008; Keinan et al. 2009; Emery et al. 2010), methods have been developed to estimate the BSR using maximum likelihood analysis of the SFS (Musharoff et al. 2019) and Bayesian analysis of multi-population genealogies (Clemente et al. 2018). The effects of the BSR we report could stimulate the development of methods that jointly consider multiple facets of variation to estimate the BSR using composite likelihood or approximate Bayesian computation. Comparison of BSR estimates across a diverse collection of populations and species could point to biological drivers of this fundamental reproductive parameter.

## Materials and Methods

### Theoretical Approach to Simulate the Breeding Sex Ratio

To rapidly simulate datasets across a range of BSR values, we followed established theory connecting BSR with effective population size (*N_e_*) (Caballero 1995; Musharoff et al. 2019). We calculated the *N_e_* of an autosomal locus as:

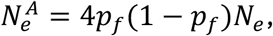

where pf is the proportion of breeding females in the population (N_f_ /N_e_). We calculated the *N*_e_ of an X-linked locus as:

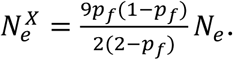

These effective population sizes were used as input to coalescent simulations.

### A Pipeline for Joint Simulation of the Breeding Sex Ratio and Demographic Change

To simulate the interaction of the BSR and demographic history, we developed a pipeline that utilizes coalescent simulations from the package *msprime* v1.2.0 (Baumdicker et al. 2022). The process began by providing values of *N*_e_, mutation rate, and recombination rate (Figure 5). We specified an *N*_e_ of 10,000, a mutation rate of 5 x 10^-8^ (per-site, per-generation), and a recombination rate of 5 x 10^-8^ (per-site, per-generation) for the results shown. After specifying these parameters, both a demographic history and a BSR were applied. Using *msprime*’s demography objects, we constructed a population bottleneck with size changes specified at defined time points. We focused our examination of demographic history on population bottlenecks because they are commonly experienced by natural populations and such changes are known to affect genomic patterns of variation.

**Figure 5.**
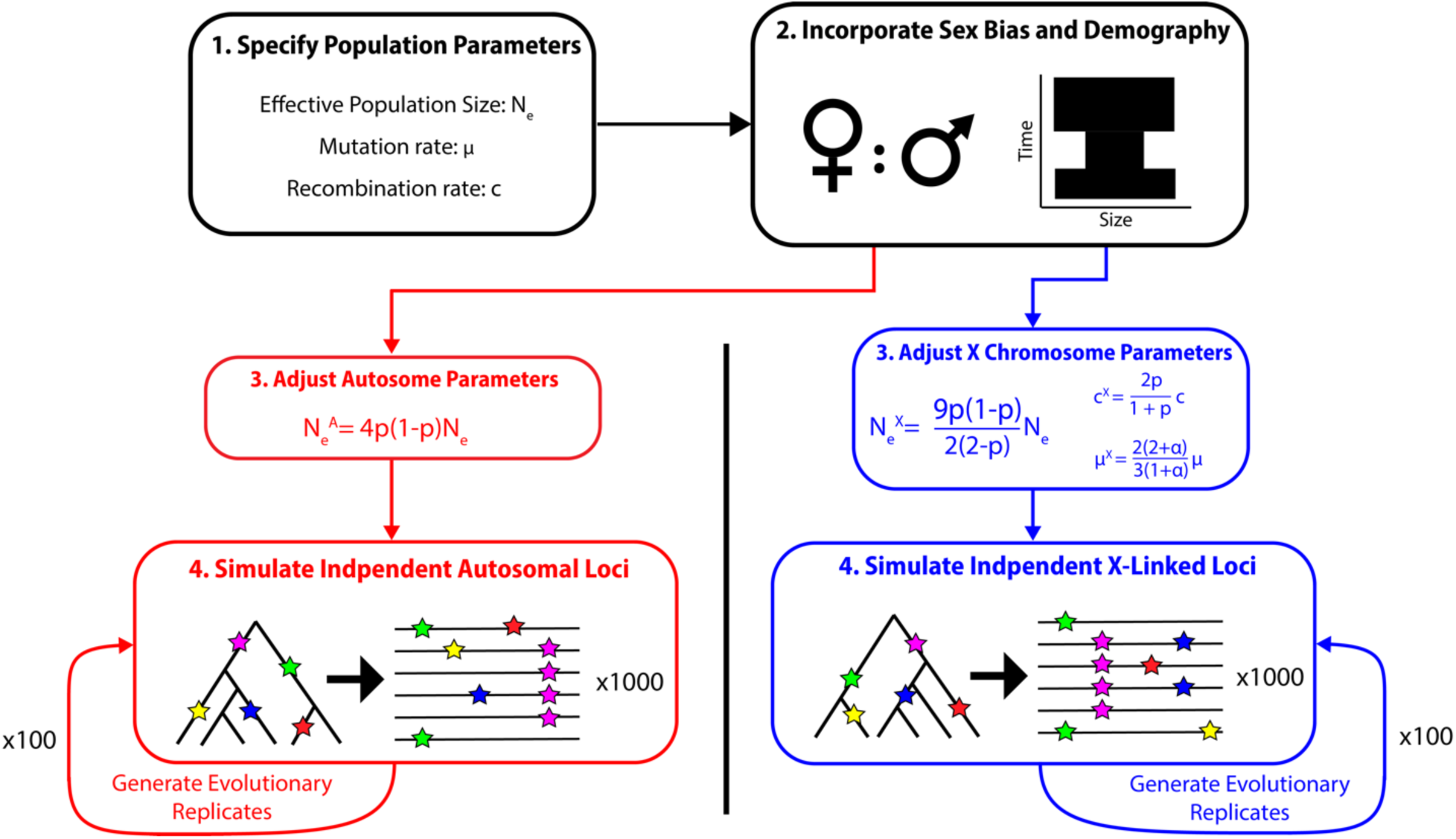
Summary of simulation framework used for generating X-linked loci and autosomal loci for comparison under specified BSRs and demographic histories.

After initializing a demographic history, we imposed a BSR on the population, using Equations 1 (autosomal loci) and 2 (X-linked loci) to adjust *N*_e_ for each epoch. To accommodate their need for separate adjustment of *N*_e_, we simulated autosomal loci and X-linked loci independently. To account for effects of the BSR on the recombination rate at X-linked loci, we rescaled the recombination rate as:

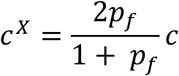

(Labuda et al. 2010; see Lohmueller et al. 2010), where *c* is the original (per-site, per-generation) recombination rate and *pf* is again the proportion of the breeding population that is female. By adjusting the recombination rate in this manner, we could capture the average behavior of X-linked loci without needing to simulate separate sexes. To account for the effects of sex differences in mutation rate, we rescaled the mutation rate at X-linked loci as:

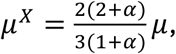

(Miyata et al. 1987), where μ is the (per-site, per-generation) mutation rate and 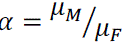 is the ratio of the male mutation rate and the female mutation rate. For autosomal loci, mutation rates and recombination rates were unchanged.

After making the adjustments described above, we performed simulations with two levels of replication with the goal of mimicking genomic patterns of variation in population samples from natural populations. We first simulated X-linked loci and autosomal loci as 10 kb windows. For each window, *msprime* generated an independent genealogy and added mutations according to input parameters. For each window, we organized *msprime* output as a dataset of single nucleotide polymorphism (SNP) genotypes in variant call format (VCF) for calculating population genetic summary statistics. All summary statistics were computed using the package *scikit-allel* v1.3.5 (https://github.com/cggh/scikit-allel). We used this process to simulate a genomic dataset composed of 1,000 X-linked loci and 1,000 autosomal loci. Genomic patterns of variation for this dataset were summarized using the mean and variance of each summary statistic taken separately across X-linked loci and across autosomal loci. For each combination of BSR and demographic parameters, we simulated 100 evolutionary replicates. We calculated the average of the mean and variance of each summary statistic across the 100 replicates, separately for X-linked loci and autosomal loci. These values were visualized, with the 2.5th and 97.5th quantiles of the distributions used as approximate confidence intervals across genomic datasets.

### Quantifying Patterns of Variation

To capture the effects of the BSR and demographic history on genomic patterns of variation, we computed a suite of summary statistics. We calculated the average pairwise number of differences (Tajima 1983) and Watterson’s θ to quantify levels of variation. We used Tajima’s D (Tajima 1989) as a summary of the site frequency spectrum of polymorphisms. We tabulated the number of unique haplotypes and the frequency of the most common haplotype as metrics of haplotype structure. We measured average pairwise r^2^ (after removing singleton SNPs) (Hill and Robertson 1968) to describe linkage disequilibrium. We treated phase as known.

### Simulated Parameter Values

To determine how relationships between the BSR patterns of variation are affected by demographic history, we simulated a range of population bottlenecks. All scenarios were simulated across a full range of BSR, starting with a proportion of females in the population equal to 0.1 (10% female, 90% male) and increasing by 0.05 until ending at 0.9 (90% female, 10% male). Each bottleneck was modeled as an instantaneous reduction in *N_e_*, a period at reduced *N_e_*, and an instantaneous increase to return to the original *N_e_*. For each of three bottleneck parameters – strength, timing, and duration – we specified three values. For bottleneck strength, we simulated 50%, 80%, and 95% reductions in *N_e_*. For bottleneck timing, we simulated a size recovery occurring 500, 1000, and 5000 generations ago. For bottleneck duration, we simulated bottlenecks lasting 100, 500, and 1000 generations. We simulated every combination of parameters, for a total of 27 unique bottlenecks.

We also considered how mutation rate and recombination rate affect the relationship between BSR and patterns of variation. We simulated the following scenarios. First, we specified mutation and recombination rates of 5 x 10^-8^ (per-site, per-generation). Second, we specified mutation rate and recombination rates of 5 x 10^-9^ (per-site, per-generation), but the results were qualitatively similar to the results we show. Finally, we considered a mutation rate three times higher in males than in females (*ɑ* = 3). We ran simulations in parallel using the Center for High Throughput Computing (CHTC) at the University of Wisconsin-Madison.

## Data Availability Statement

The datasets used to produce each figure of this study can be found in the Supplementary Material (Supplementary Tables 1 & 2). The computational scripts that were used to generate these data can be found on online at (https://github.com/PayseurLabUWMadison/BSR_simulations).

## Acknowledgements

We thank all members of the Payseur Lab for thoughtful discussions about this work. We additionally thank members of UW-Madison’s Center for High Throughput Computing for providing computational resources and helpful feedback. This work was supported by the National Institutes of Health (NIH) grant R35GM139412 (to B.A.P.). W.J.S. was partially supported by the NIH Graduate Training Grant in Genetics at the University of Wisconsin-Madison (T32GM007133).

## Author Contributions

W.J.S. and B.A.P. both designed this study and wrote the manuscript. W.J.S. performed the research with supervision from B.A.P.

