## Supplementary Figures for "Comparative Patterns of Variation on the X Chromosome and Autosomes: The Role of the Breeding Sex Ratio"

#### Supplementary Material

Supp. Fig. 1: Effects of BSR on Inter-locus Variances in Populations of Constant Size

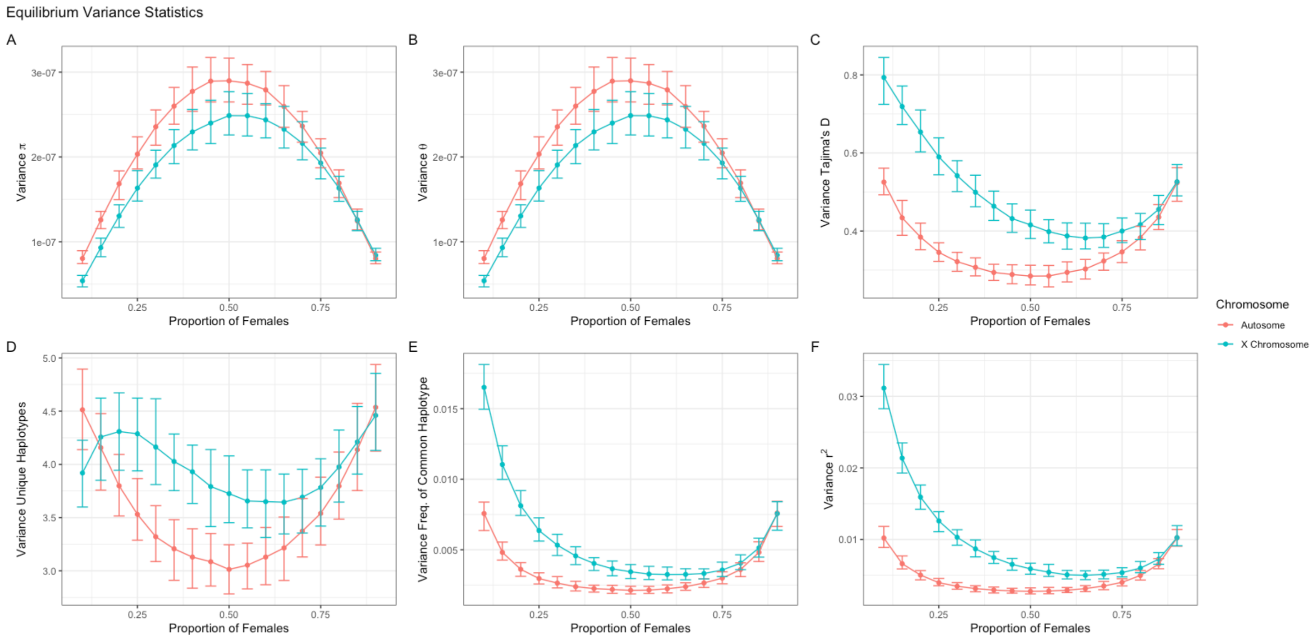

Supp. Fig. 2: Effect of Bottleneck Strength on Means across Loci: Longer, Distant Bottleneck

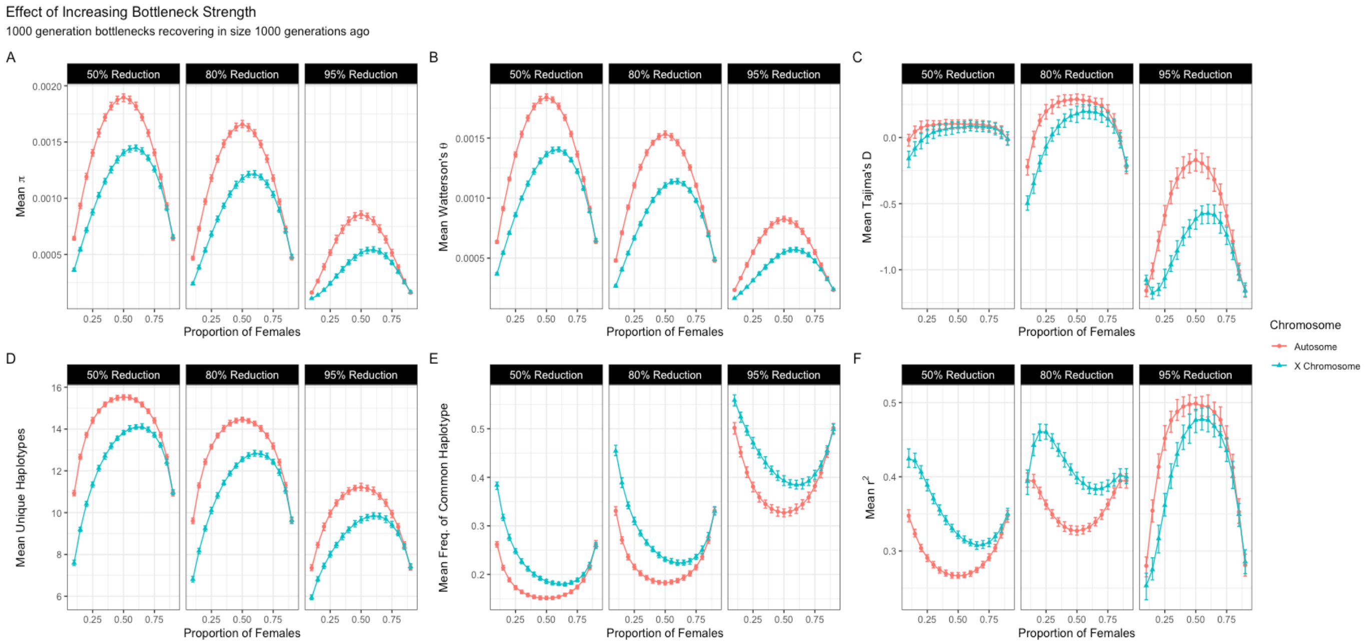

### Supp. Fig. 3: Effects of Bottleneck Strength on Means across Loci: Longer, Very Distant Bottleneck

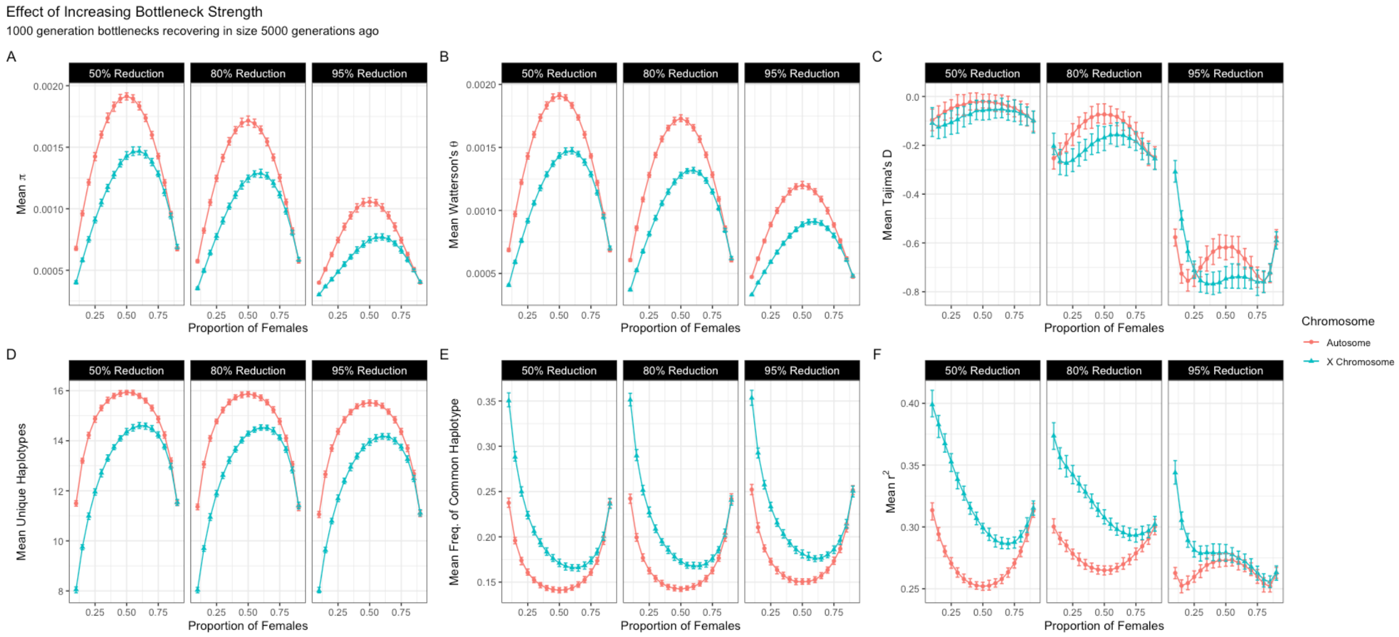

### Supp. Fig. 4: Effect of Bottleneck Strength on Inter-locus Variances

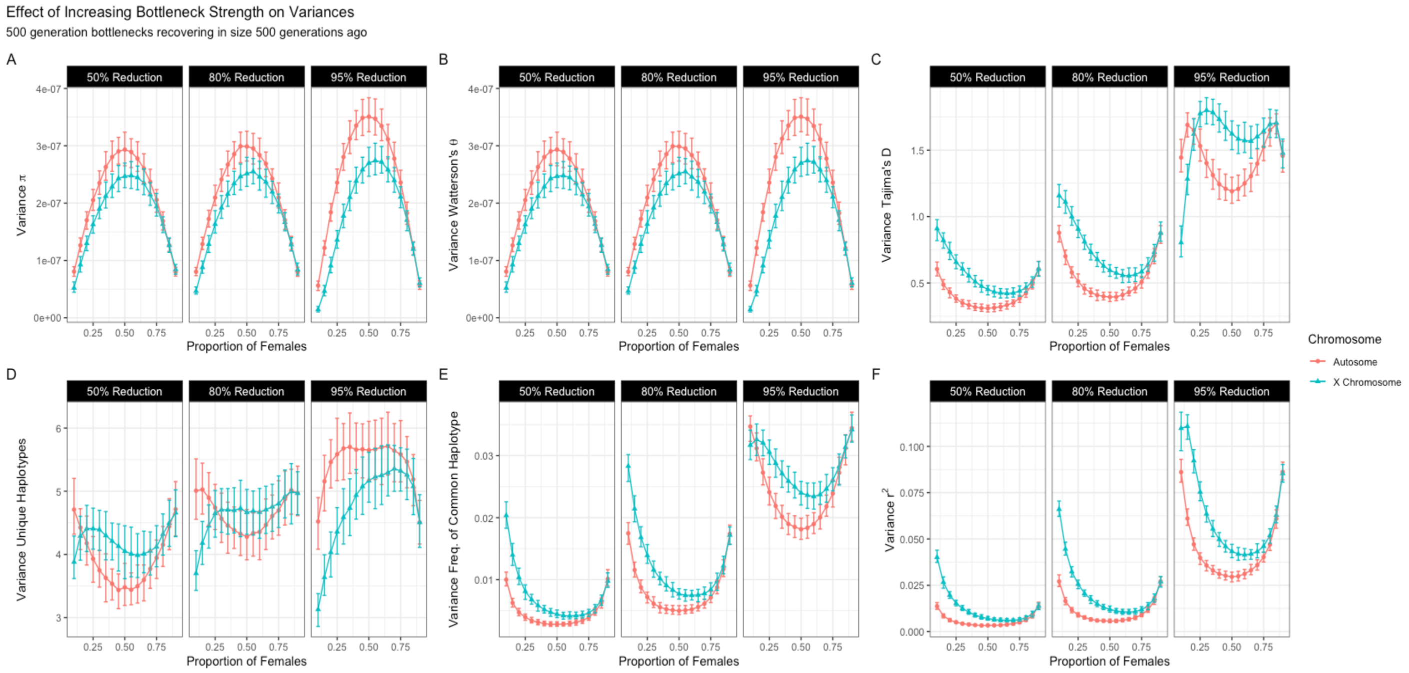

### Supp. Fig. 5: Effects of Bottleneck Duration on Means across Loci: Weaker Bottleneck

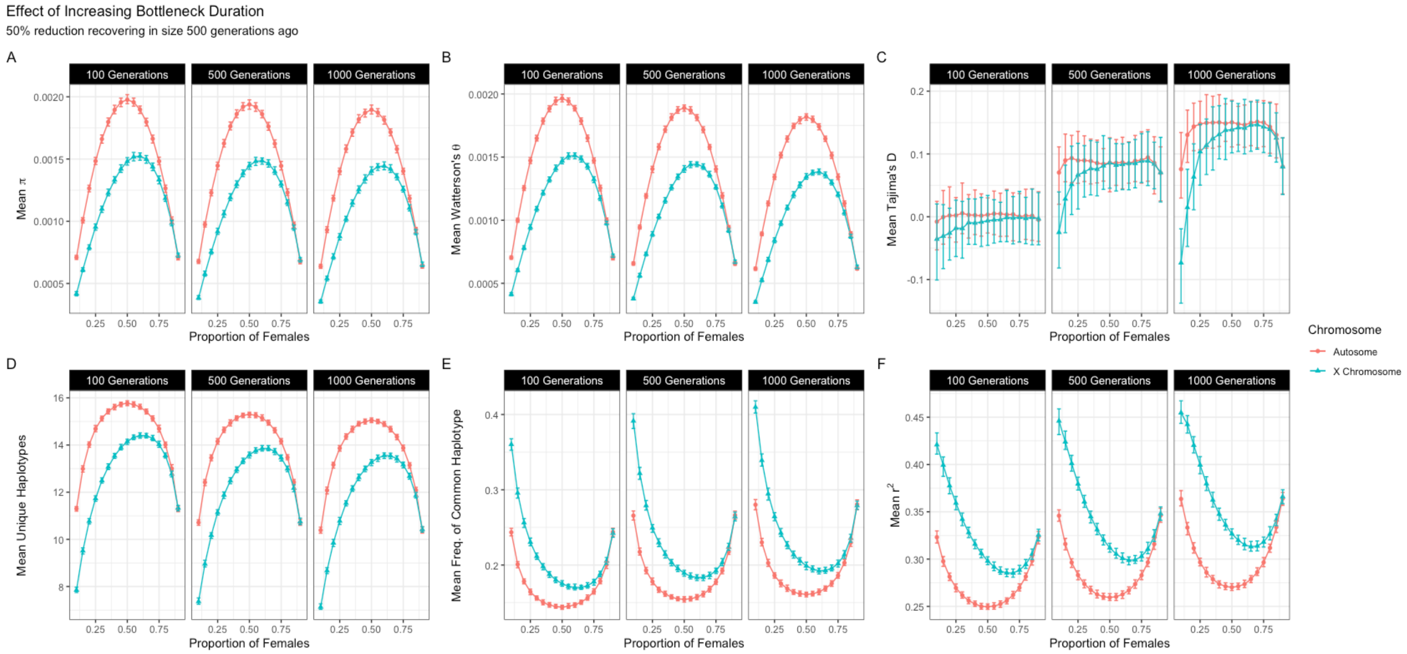

Supp. Fig. 6: Effects of Bottleneck Duration on Means across Loci: More Distant Bottleneck

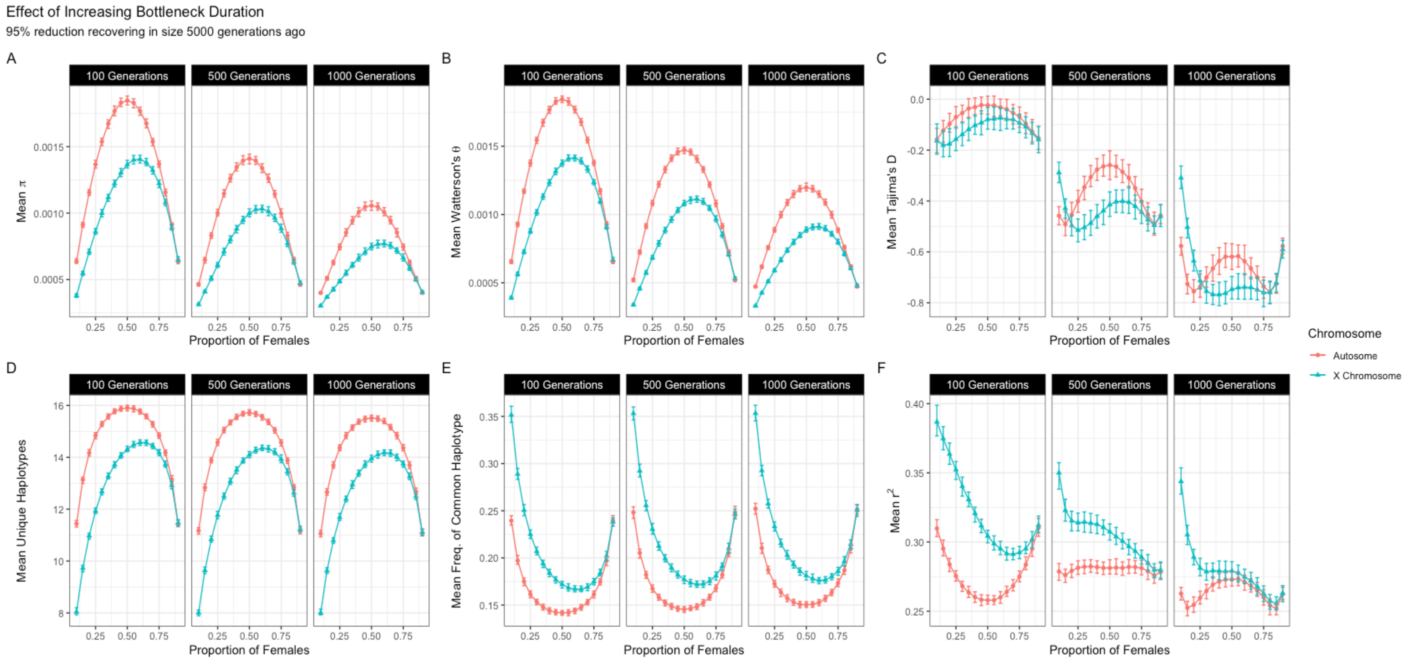

Supp. Fig. 7: Effect of Bottleneck Duration on Inter-locus Variance

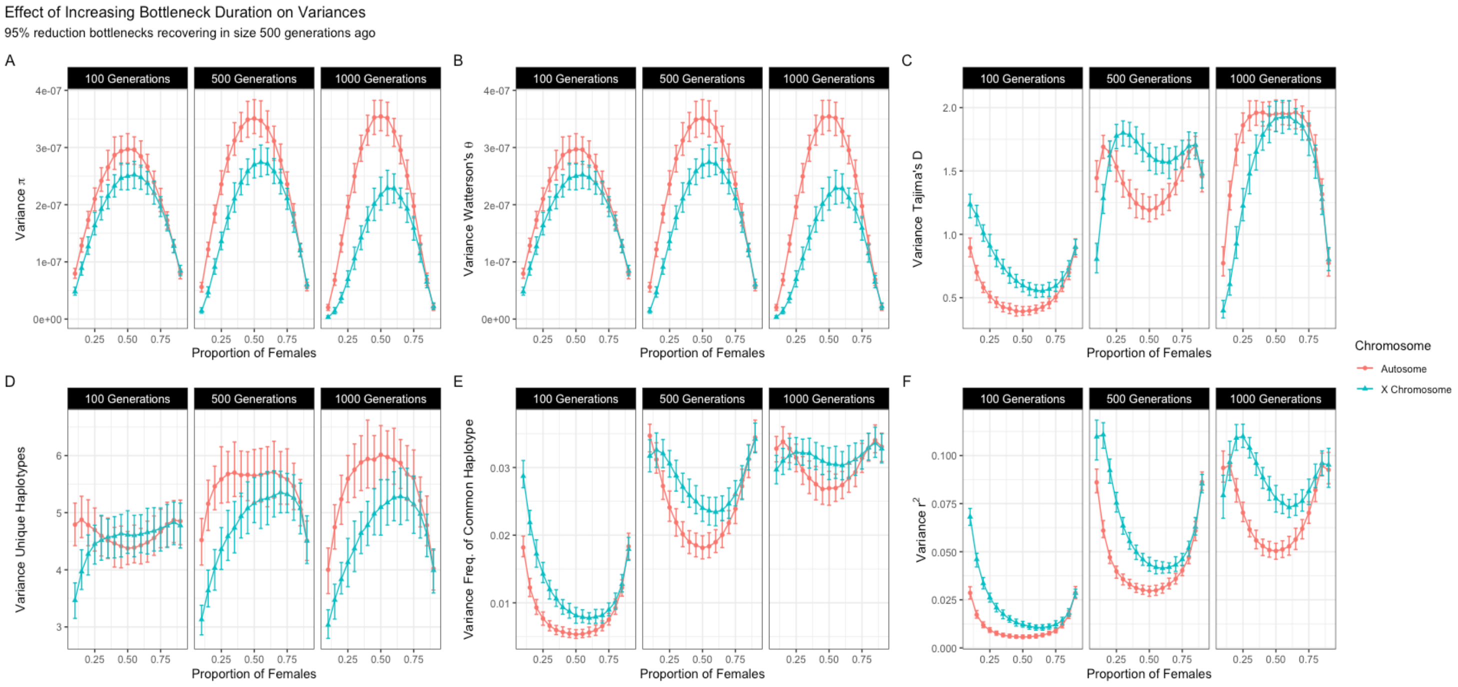

### Supp. Fig. 8: Effects of Bottleneck Timing on Means across Loci: Weak Bottleneck

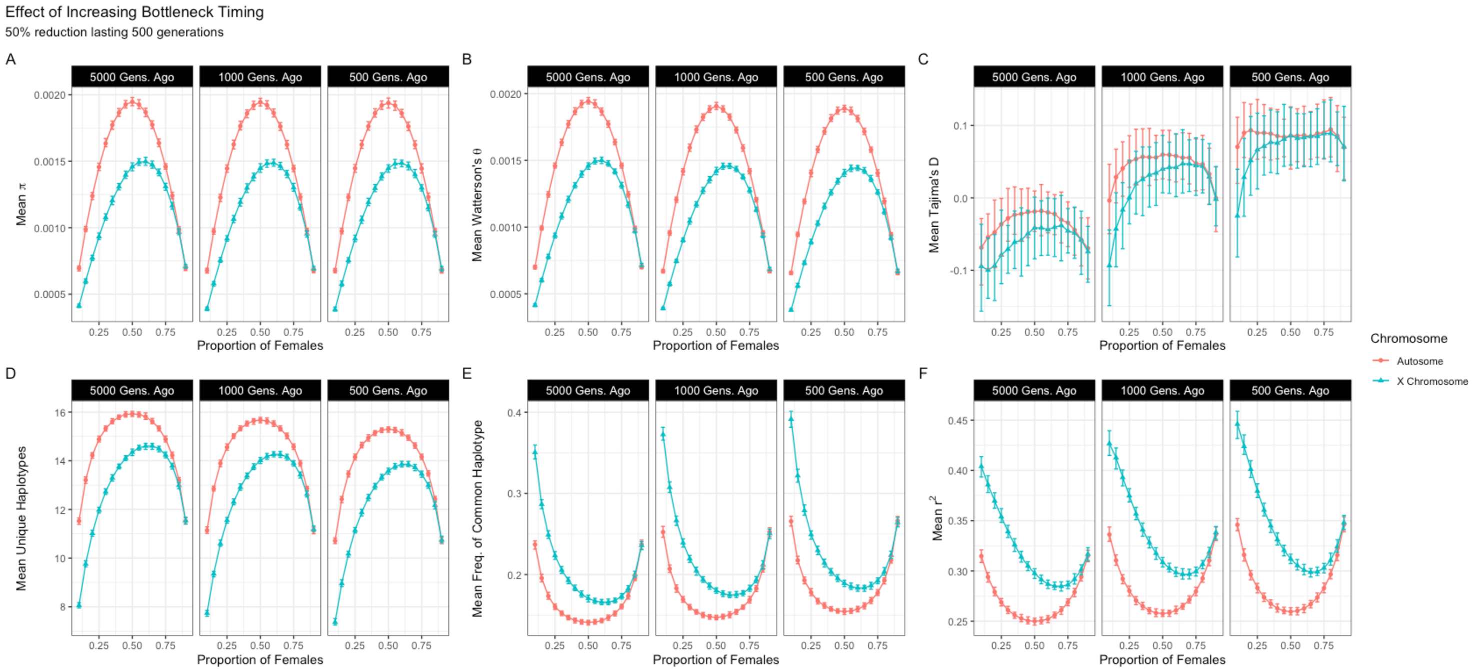

### Supp. Fig. 9: Effects of Bottleneck Timing on Inter-locus Variances

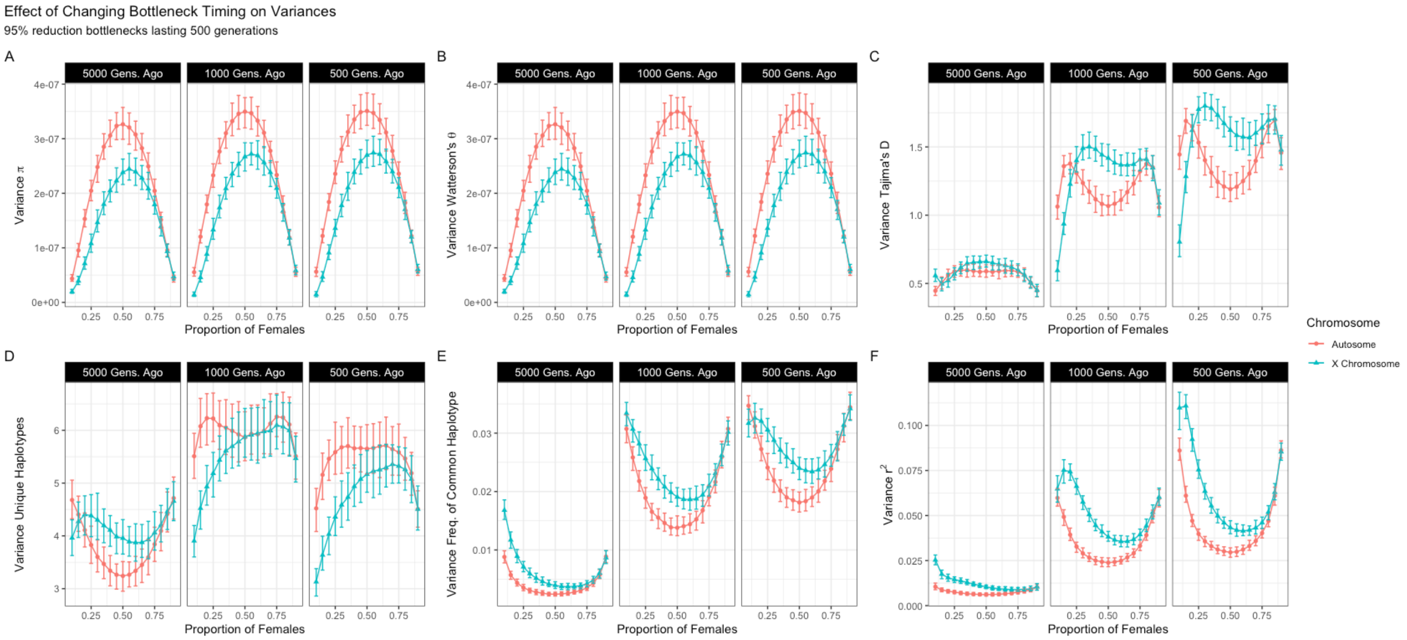

Supp. Fig. 10: Average SFS of Autosomal and X-linked Loci at Constant Size

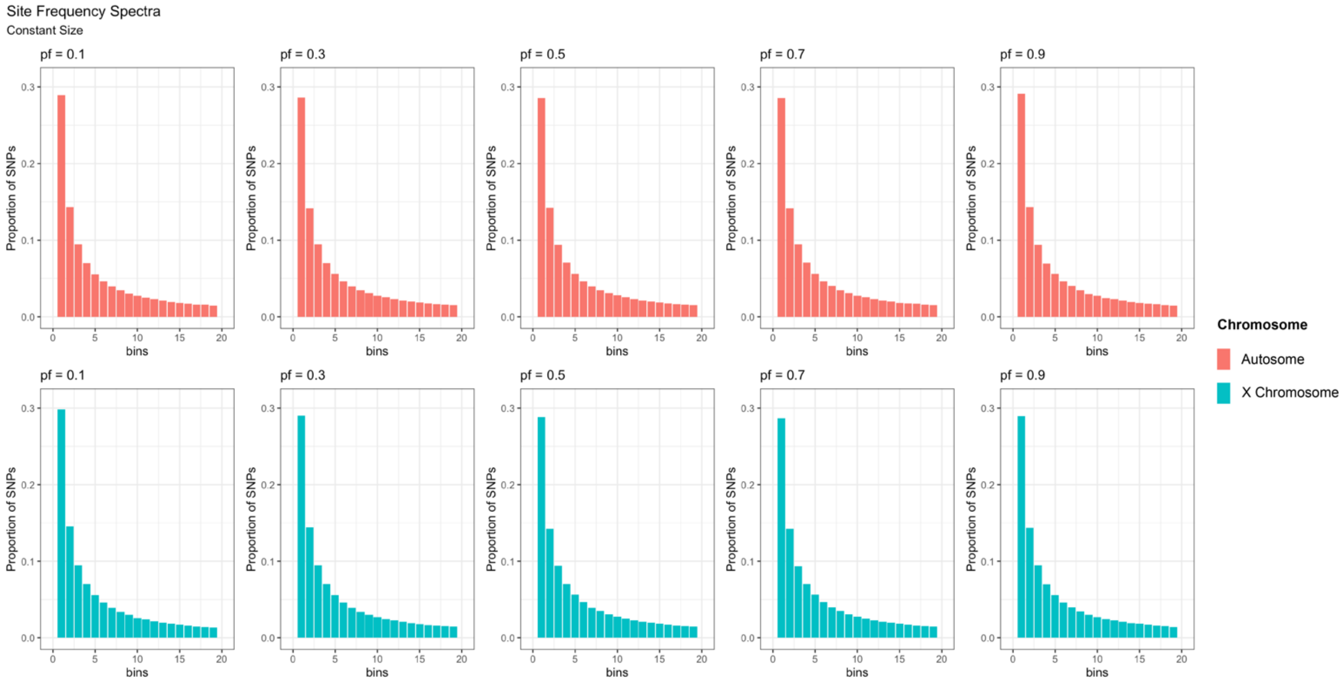

Supp Fig. 11: Average SFS of Autosomal and X-linked Loci Under Bottleneck

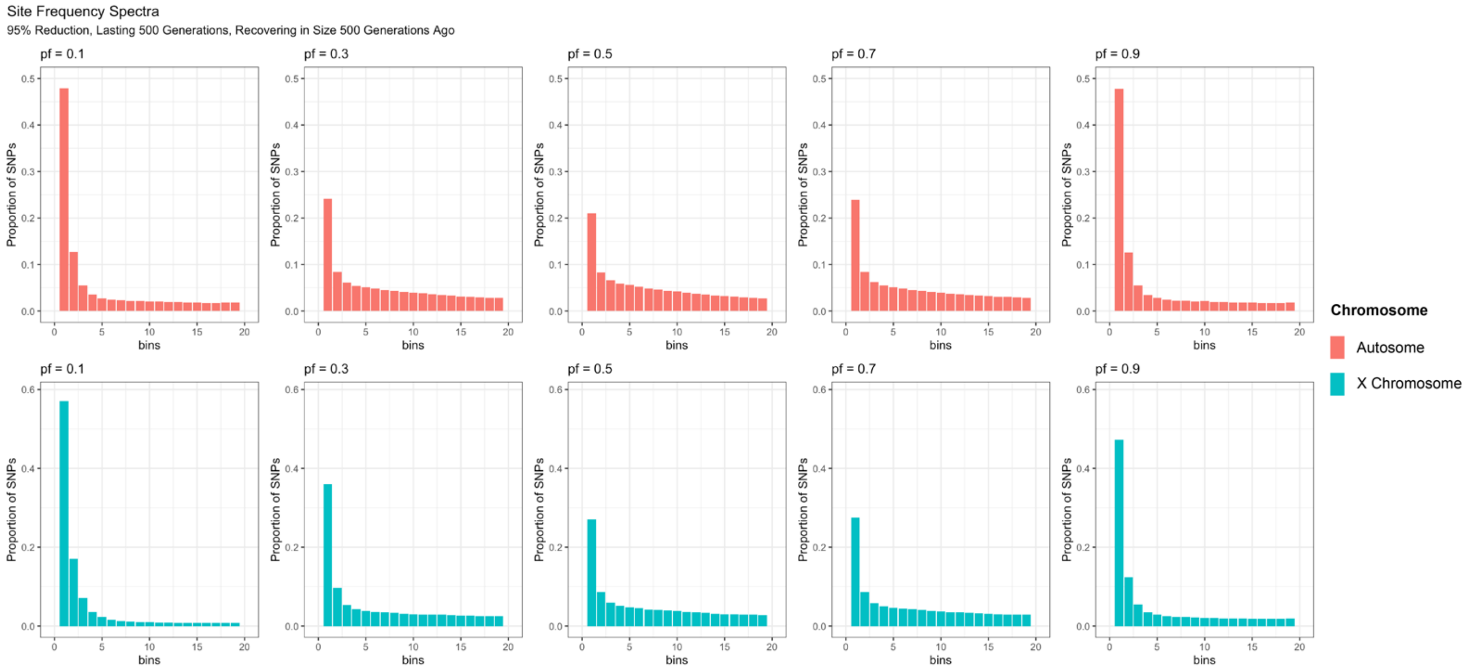

Supp. Fig. 12: TMRCA of $r^2$ Distribution

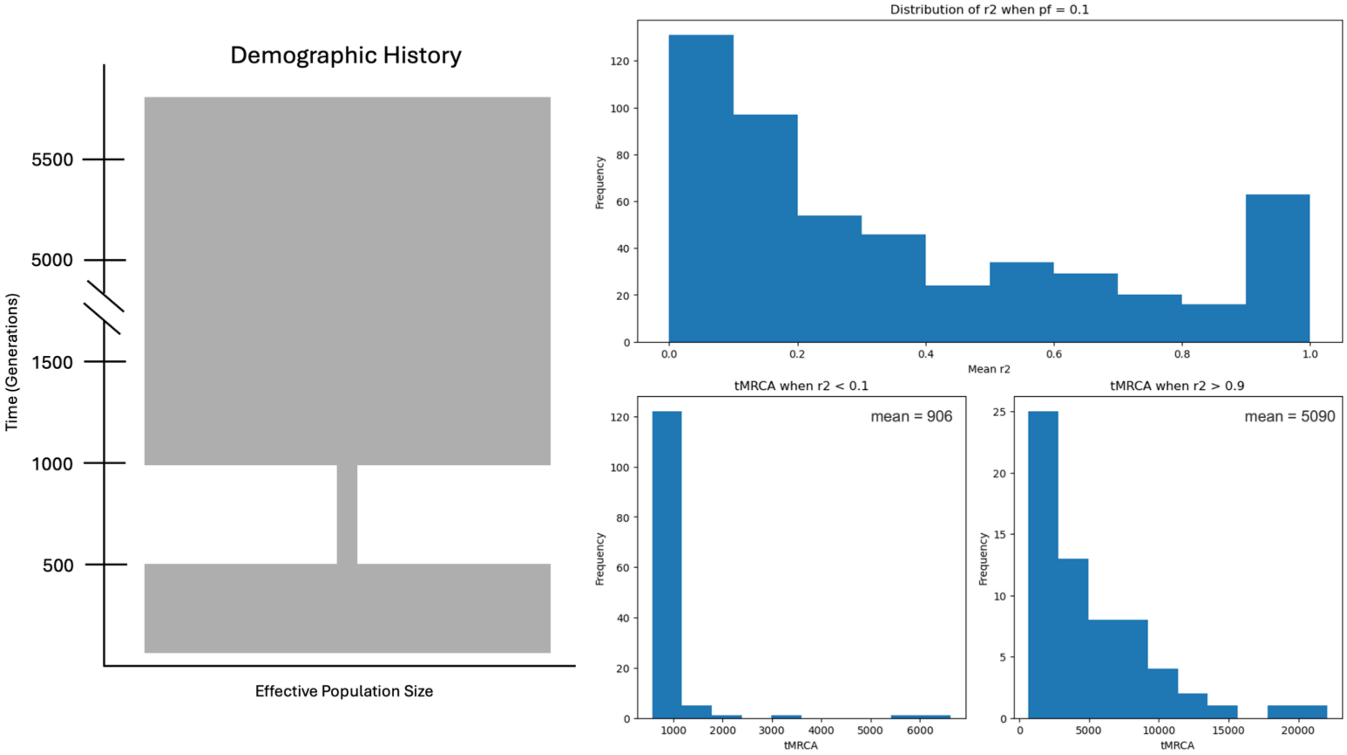

Supp. Fig. 13: Effects of Male-Biased Mutation Rate Means Across Loci: Constant Size

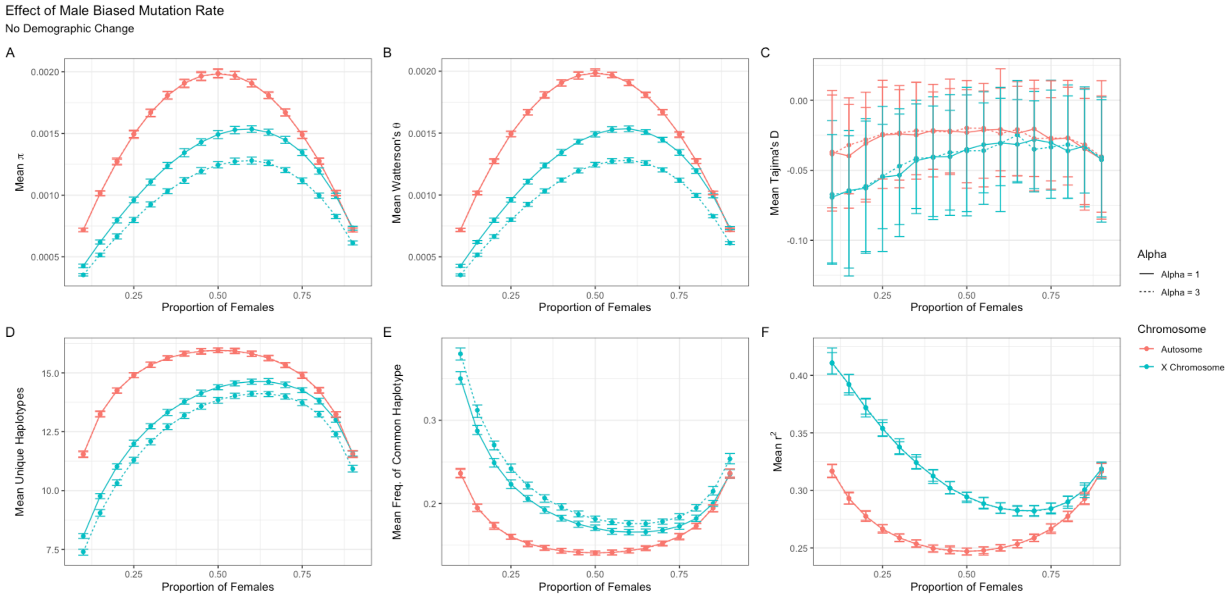

Supplementary Figure 14: Effects of Male-Biased Mutation Rate Means Across Loci: Bottleneck

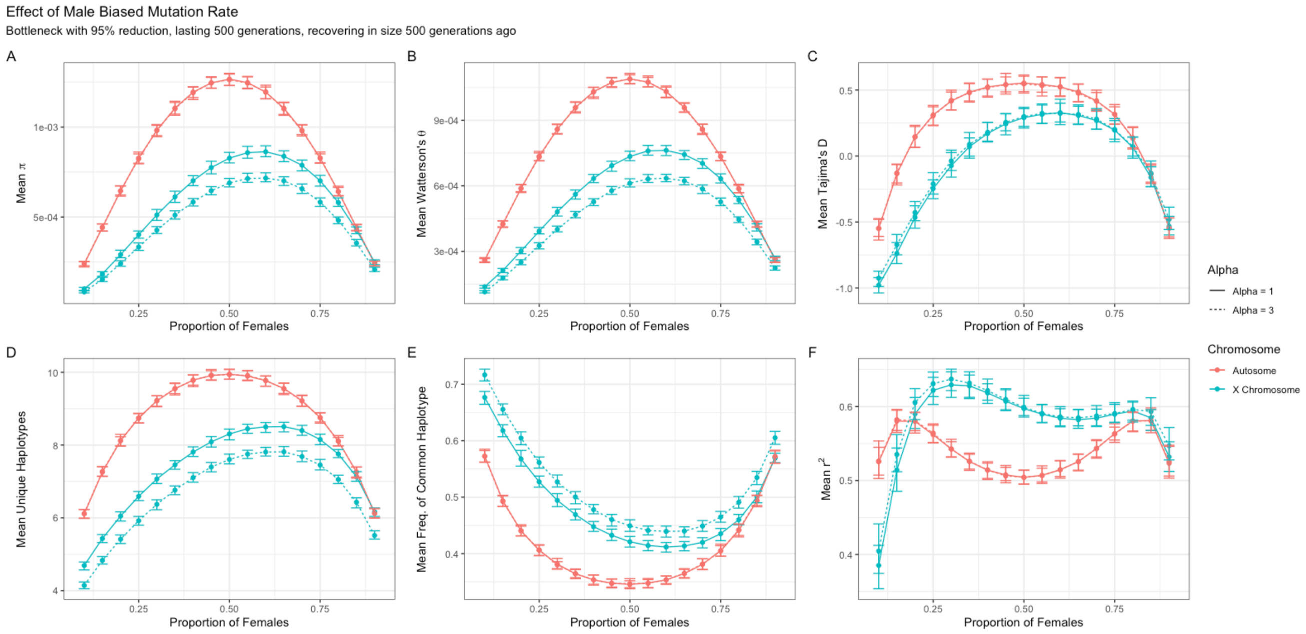
